# Understanding land-use response of bird communities in an arid ecosystem of India

**DOI:** 10.1101/2020.09.27.315473

**Authors:** K Varun, Sutirtha Dutta

## Abstract

a. The Indian Thar desert has lost much of its grasslands over the last few decades, mainly due to land-use change from pastoralism to agriculture. Expanding croplands and intensifying grazing pressures are popularly hypothesized to be major drivers of biodiversity loss in the region. Our study aims to investigate the effects of contemporary land-use change on bird communities of the Western Thar Desert.
b. We surveyed 58 randomly laid line transects in a c2000 sq.km study area, to quantify parameters of bird community structure in three predominant land-use types viz. protected grasslands, rangelands, and non-irrigated croplands. Fieldwork for the study was conducted in the dry season (winter and summer) between December 2018 and April 2019.
c. During winter, overall bird richness and abundance were highest in protected grasslands followed by non-irrigated croplands and rangelands. Protected grasslands also had a higher abundance of diet and habitat specialists. Compared to protected grasslands, density was lower in non-irrigated croplands and rangelands for 35% and 10% of species, respectively. A majority of the negatively affected species were insectivorous grassland specialists.
d. Contrary to the pattern in winter, overall bird richness, abundance, community composition, and guild structure in summer were similar across three land-use types. Only one of the 17 analysed species had lower density in modified land-use types.
e. Overall, protected grassland was the best habitat for birds and was specifically important for specialists, particularly during the winter. Rangelands and fallow croplands sustained most generalists at comparable densities but had severe negative impacts on specialists.
f. Synthesis and application: Our results point out that low-intensity agro-pastoral land-uses can supplement, but not replace, protected areas in conservation of Thar desert’s avifaunal diversity. Our results are consistent with the idea of managing dryland habitats as agro-grassland mosaics with embedded protected areas, in order to reconcile human needs and biodiversity conservation at a landscape scale.

## 1. Introduction

Drylands constitute about 41% of the world’s surface and are home to a large spectrum of biodiversity (Reynolds et al., 2007). They also support over two billion people who are predominantly poor and depend on the land for their food and livelihoods (Durant et al., 2015; Reynolds et al., 2007). Historically, most drylands were home to (semi) nomadic pastoralists who used grassland-savannah habitats for free-range livestock grazing. Cultivation was secondary and limited to seasonal farming of a few hardy crops that could tolerate the low and erratic rainfalls. Such low-intensity human use prevailed over historical times in many drylands, allowing some wildlife to adapt to, and even benefit from such habitat modifications (Safriel et al., 2010). However, in the last few decades, advanced irrigation facilities coupled with open market policies have propelled the expansion of crop-based agriculture over traditional rangelands (Safriel et al., 2010). Simultaneously, grazing pressures on many rangelands have increased, partly due to agricultural diversion of pastures, often leading to the degradation of semi-natural habitats (Jeuken, van den Berg, Alkemade, de Leeuw, & Reid, 2012). Expansion and intensification of farming and livestock grazing are widely considered to be the biggest threats to arid and semi-arid drylands of the world (Safriel et al., 2010). Yet, their impacts on biodiversity are poorly studied; especially in the tropical developing world, where most drylands are located, and the scale and magnitude of land-use activities are the highest.

Diversion of natural land for agriculture is known to have a wide range of ecological impacts on biodiversity (Foley et al., 2005; Newbold et al., 2015); and birds are excellent bioassays for studying such impacts (Green, Cornell, Scharlemann, & Balmford, 2005; Sutherland, 2004). Land-use response studies on birds, majorly conducted in various forested ecosystems of the world, have shown that progressive change in land-use generally causes – a) decrease in species richness and abundance of birds at both local and landscape scale (Newbold et al., 2015); b) precipitous decline in abundance of specialized species such as understory insectivores, and other sensitive groups (Newbold et al., 2012); c) a significant change in community structure beyond a certain threshold (Elsen, Ramesh, & Wilcove, 2018). Notwithstanding this general pattern, many low-intensity land-uses also provide important secondary habitats for some forest birds (Bhagwat, Willis, Birks, & Whittaker, 2008; Elsen et al., 2018; Hendershot et al., 2020; Hughes, Daily, & Ehrlich, 2002; Ranganathan, Daniels, Chandran, Ehrlich, & Daily, 2008). Contrary to forested systems, effects of agriculture on dryland birds are much more nuanced; and contingent upon species’ ecology, region, season, land-use intensity, etc (Kamp, Urazaliev, Donald, & Hölzel, 2011; Rodríguez-Estrella, 2007; Wolff, Paul, Martin, & Bretagnolle, 2001; Wright et al., 2012). Limited knowledge from dryland systems suggests that, on one hand, low-intensity farming and livestock grazing supports, and even benefits, many open-habitat birds (Kamp et al., 2015; Wolff et al., 2001; Wright et al., 2012). On the other hand, it threatens a few other species that show irreplaceable dependency on primary or near-natural habitats (Brennan & Kulvesky, 2005; Rodríguez-Estrella, 2007). To add to the complexity, some species utilize modified land-uses during the resource-scarce dry season but are dependent on near-natural habitats during the breeding season (Dutta, 2012). Given this complexity, meaningful inferences and generalizations at larger scales require detailed information on the land-use response from several study taxa and systems (Laurance et al., 2014).

With human population and food demands constantly rising, biodiversity conservation increasingly relies on multiple-use areas such as croplands and rangelands (Barlow et al., 2018; Kremen & Merenlender, 2018; Laurance, Sayer, & Cassman, 2014; Tilman, Balzer, Hill, & Befort, 2011). This dependency is even stronger in arid-semiarid landscapes of developing countries, where protected areas are inadequate and wildlife shares space with economically deprived people who rely on subsistence agro-pastoral livelihoods (Dutta & Jhala, 2014; Singh et al., 2006). This interface necessitates identifying land-use practices beneficial to biodiversity so that mechanisms can be developed to maintain them before they are irreversibly lost (Wright, Lake, & Dolman, 2012).

### 1.1 Land-use change in the Thar desert

The Thar Desert, lying in the northwestern corner of India, is the world’s most populated desert and supports many bird species including the globally threatened Great Indian Bustard *Ardeotis nigriceps* (Singh et al., 2006). Historically, this area was sparsely populated with many nomadic or semi-nomadic pastoral communities, and a few settlements, with agriculture, restricted to low-lying areas. Since the 1980s, various irrigation schemes including the flagship Indira Gandhi Canal promoted cultivation in vast stretches of the desert (A. R. Rahmani & Soni, 1997) and led to an immigrant population boom with the number of people increasing almost tenfold in the next 40 years (Dhir, Joshi, & Kathju, 2018). Many local pastoral societies became sedentary and started cultivation along with animal husbandry. Consequently, grasslands and rangelands were converted to croplands at a massive scale (Tian, Banger, Bo, & Dadhwal, 2014). Livestock populations also increased simultaneously, benefiting from the booming dairy industry (Dutta, 2018), and increased grazing pressure on remnant rangelands. Because of these developments, arid grasslands with low grazing pressures were decimated from most areas, except for the most remote western parts of the Thar desert such as the Indo-Pakistan border, Desert National Park Wildlife Sanctary, and Pokhran Field Firing Range (Islam & Rahmani, 2011). Even these areas are now experiencing agricultural intensification due to increased demand for a desert crop Guar (*Cyamopsis tetragonoloba*) that has found recent industrial use in hydraulic fracturing (Anoop, Bharadwaj, & Shekhawat, 2017). These changes are expected to adversely affect the biodiversity of the region.

The western part of the Indian Thar desert is at the junction of the extensive arid tract running from Northern Africa through the middle-east and the semi-arid tract running across western India. It is the meeting point and range limit for avifauna with diverse affinities. It also falls on the central asian flyway, acts as an important migratory ground for many passerines, and is enlisted as an Important Bird Area (Birdlife International, 2020). Considering the importance of this region for the conservation of avifauna and to bridge the knowledge gap concerning the land-use response of dryland birds, we conducted extensive bird surveys across three land-use types predominant in the western Thar desert. Our overarching aim was to understand how birds respond to various land-use regimes in the region. More specifically, we investigated:

1. How different land-use types compare in terms of their bird community structure? To answer this we looked at the following parameters: regional species pool, local species richness, bird abundance, and community composition.
2. Are responses to land-use change contingent on species ecology? Conversely, how do different ecological groups respond to individual land-use changes? Here, we divided the species assemblages based on feeding and habitat preferences of species and then compared the abundances of each group across different land-cover types.
3. What proportion of species win (benefit) and lose (decline) in response to a land-use change? To understand this, we examined how densities of individual species differ across three land-use types.

## 2. Study area and description of land-use types

The study was conducted over a 2016 sq.km (27.42 to 26.56 N, 70.89 to 70.14 E) area in the western Thar desert, administratively falling under the Jaisalmer district of Rajasthan state in northwestern India (Figure 1). The region falls under the Desert Biogeographic Zone (Rodgers and Panwar, 1988), characterized by severe summers (>45 C) and winters (<5 C) with high diurnal temperature ranges, and sparse erratic rainfall (∼200 mm). The southern part of the study area encompassed some parts of the Desert National Park Wildlife Sanctuary (hereafter, DNP WLS). The DNP WLS, notified in the early 1980s to preserve the wildlife and traditional lifestyle, is the only large protected area in the Indian Thar desert (Anoop et al., 2017; Singh et al., 2006). However, this 3000 sq.km area includes about a hundred legal settlements, and over sixty thousand people almost entirely dependent on the sanctuary for their livelihood (Anoop et al., 2017). Semi natural protected grasslands exist only in few small parts of the sanctuary (approx. 200 sq.km) where fenced enclosures have been created to regulate livestock grazing (Anoop et al., 2017; Dutta et al., 2017). Majority of the western Thar landscape outside these enclosures is composed of rain-fed croplands which are left fallow after the four-month cropping season; and semi-wild rangelands that are used for free-ranging livestock grazing throughout the year. Large parts of the study area overlapped with the priority Great Indian Bustard landscape (Wildlife Institute of India, 2020).

**Figure 1.**
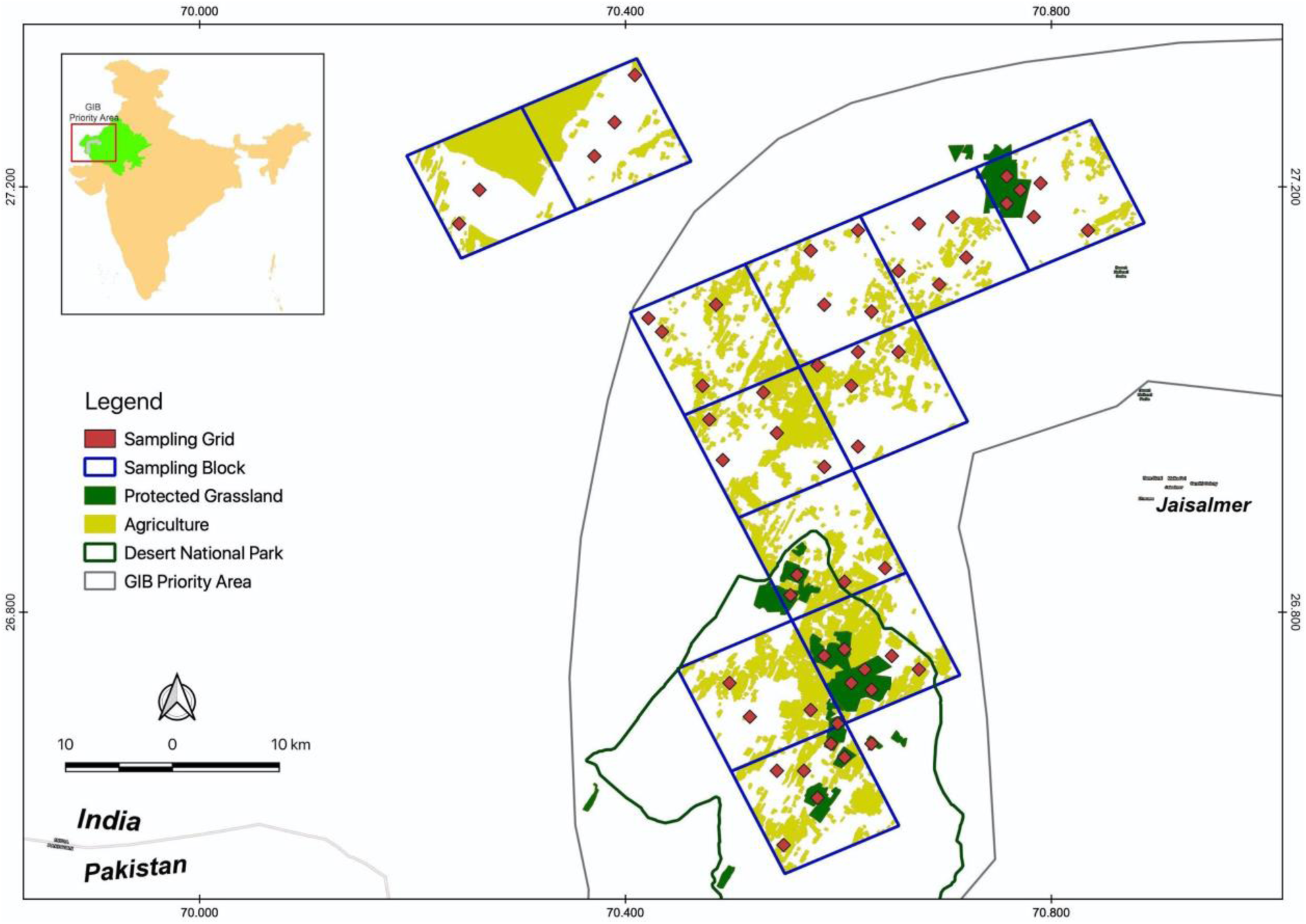
Study area map showing the distribution of transects and the three land-use types. Layers for land-use classification, GIB priority area, desert national park, protected grasslands and sampling blocks were obtained from CAMPA-Bustard Recovery program, Wildlife Institute of India (Dutta et al., 2017).

The study area was classified into three dominant land-use categories based on an ensemble of mapping techniques. Each category is described in detail below:.

1. Protected grasslands: confined to the graze-proof enclosures built up by the DNP WLS and Desert Development Program of the Forest Department. Grazing was ‘regulated’ to a minimum inside these areas and agriculture was strictly prohibited. The grazing pressure in these areas was at least an order lower than rangelands despite the prevailing drought conditions.
2. Rangelands or grazing lands: areas primarily used for grazing domestic livestock – goats, sheep, cows, and camels. They did not have any recent history of farming. These areas differed from the protected grasslands due to their high grazing pressure and human disturbance.
3. Extensive croplands (non-irrigated and rainfed or fallow croplands): most parts of the Thar desert are not irrigated and receive erratic rainfall, thus limiting farming to monsoon or the ‘Kharif’ cropping season, that too in years of good rainfall. The land is left fallow after cropping and allowed to regenerate through dry months. All non-intensive/rainfed croplands in the study area were less than two-year-old regenerating fallows at the time of sampling. Mostly, the last crop cultivated on these non-irrigated farmlands was Guar (*Cyamopsis tetragonoloba)*.

## 3. Methods

### 3.1 Study design and field methods

We placed 59 line transects of 1km length randomly across the study area, in proportion to the area in each land-use category. This approach of stratified random sampling resulted in 15 transects in protected grasslands, 26 transects in non-irrigated croplands, and 18 in rangelands. All transects were oriented in a north-south direction and were placed at least 1 km apart from each other to maintain spatial independence. Bird surveys were conducted in the dry season of 2018-19 – winter (December 25 to February 20), when the winter migrants were present, and early summer (21 March to 21 April), when many resident species start territory formation. All transects were surveyed three times in a season to reduce the noise induced by temporal unavailability and non-detection of species, and thereby achieve accurate and precise sample estimates of local bird populations. Each transect was walked at a slow uniform pace and birds were scanned using a 8×42 Nikon binocular. Each bird seen or heard on the transect was identified and recorded in a conventional distance sampling protocol (Buckland, Rexstad, Marques, & Oedekoven, 2015).

### 3.2 Analytical Methods

#### 3.2.1 Comparative species richness, abundance, and guild structure at sites

We modelled species richness at every transect as a function of land-use type (categorical variable) using Generalised Linear Models (GLM) of the poisson family in a hypothesis-testing framework. We used average species richness of protected grasslands as our control and compared average species richness of all other land-use types against it, segregating the analyses by season.

Site-level abundance was confounded with variability in detection of species. Hence, we corrected raw counts of birds by modelling species-specific detection functions in a Distance sampling framework (Buckland et al., 2015), while also testing for the effect of land-use on the detection of individual species. As many species had inadequate records for modelling detection, we clubbed similar species together based on their taxonomic affinity, appearance, perch preference, etc. (Supplementary information). The corrected counts were then standardized to one sq.km to obtain density (abundance per sq.km) of all species for a given plot. To check how species’ density varied across different land-use types, we used general linear model (LM) in a hypothesis-testing framework and modelled the non-detection corrected log-transformed density as a function of categorical land-cover types. The null hypothesis for this test was that mean density in a particular land-use was equal to the mean density in protected grasslands (control).

To evaluate the guild structure of the community, all species were classified into five feeding guilds viz. carnivores, granivores, insectivores, nectarivores, and omnivores, following the Elton traits database (Wilman et al., 2014). A species was assigned to one of the four specialist feeding guilds (carnivore, granivore, insectivore, or nectarivore) if more than 60% of its diet was constituted of one food resource (meat, plant-seed, invertebrate, nectar respectively). However, as carnivorous and nectarivorous species were represented by very few species (n=1), we omitted them from feeding-guild specific analyses. Species were separately classified into three categories – grassland/desert specialists, habitat generalists, and others – based on their broad habitat preferences. Habitat classification was done by reviewing relevant published literature and prominent field guides (BirdLife International, 2020; Billerman et al., 2020; Grimmet, Inskipp & Inskipp, 2011; Ali & Ripley, 1998). Since forest and wetland specialists were not represented by many species, they were clubbed together as others during analysis.

#### 3.2.2 Regional species pool and local community composition

Adequacy of sampling, in terms of capturing all species present in a land-use type, was assessed by plotting accumulation and rarefaction curves of species against sites (Gotelli & Colwell, 2001). Estimates of landscape-scale species richness were obtained using the bootstrap method. To visualise how bird communities are structured in different land-use types, we conducted Non-metric multidimensional scaling (NMDS) ordination on an abundance-based community matrix with Bray-Curtis index of (dis)similarity. To check the statistical significance of differences in community composition between land-use types, we performed a pairwise permutational multivariate analysis of variance (perMANOVA) on the same dataset (Anderson, 2017). perMANOVA checks if groups are significantly different from each other in a multidimensional space. We further evaluated the average magnitude of compositional difference between bird communities using multivariate Analysis of similarity (ANOSIM).

#### 3.2.3 Species level trends

To evaluate species-specific trends in response to land-use, we compared the observed encounter rate of each species between different land-cover types with protected grasslands as control.

Observed encounter rates of species were used as a proxy for actual density as detection probability did not vary significantly with land-cover type for any species. The log-transformed encounter rate of a species was modelled as a linear function of land-cover with protected grasslands as control (intercept). The resulting species-specific response was then classified into different trend categories based on model estimated effect size and its confidence interval. The difference was considered to be statistically and biologically significant if the confidence intervals of encounter rate for any land-use type did not overlap that of Protected grassland at alpha=0.10 and the difference in their mean values (or the effect size) was greater than 20%. A detailed summary of species-specific parameters is provided in the supplementary material. Species with few sightings (less than 10 per season) or low encounter rate (less than 0.1 individual/km) were omitted from this analysis.

All the analyses were done on the R statistical platform and the source code is provided in the supplementary information.

## 4. Results

### 4.1 Local Species Richness and bird abundance

#### Winter

Models with land-cover type as a predictor of species richness and abundance did not explain the data better than null models (deltaAICc > 4), indicating that land-use did not influence diversity. When compared between land-cover types, mean species richness in protected grasslands (Mean ± SE = 09.46 ± 0.79 [8.033, 11.156]) was only marginally higher than that in rangelands (Mean ± SE = 07.72 ± 0.65 [6.514, 9.116]; GLM: log (Difference) = −0.2 [−0.43, 0.03]) and non-irrigated croplands (Mean ± SE = 08.23 ± 0.56 [7.2, 9.409]; GLM: log (Difference) = −0.14 [−0.35,0.07]) (Figure 2.a.). However, contrary to species richness, mean abundance in protected grasslands (Mean ± SE = 454.38 ± 70.07 [336.6, 613.8]) was much higher than non-irrigated croplands (Mean ± SE = 312.2 ± 63.97 [209.8, 464.8]; LM: Difference = −198.88 [−375.14, −22.60], p-value=0.02) and rangelands (Mean ± SE = 255.70 ± 53.22 [170.8, 332.8]; LM: Difference = −142.33 [−332.39, 47.72], p-value=0.13) (Figure 2.b.).

**Figure 2.**
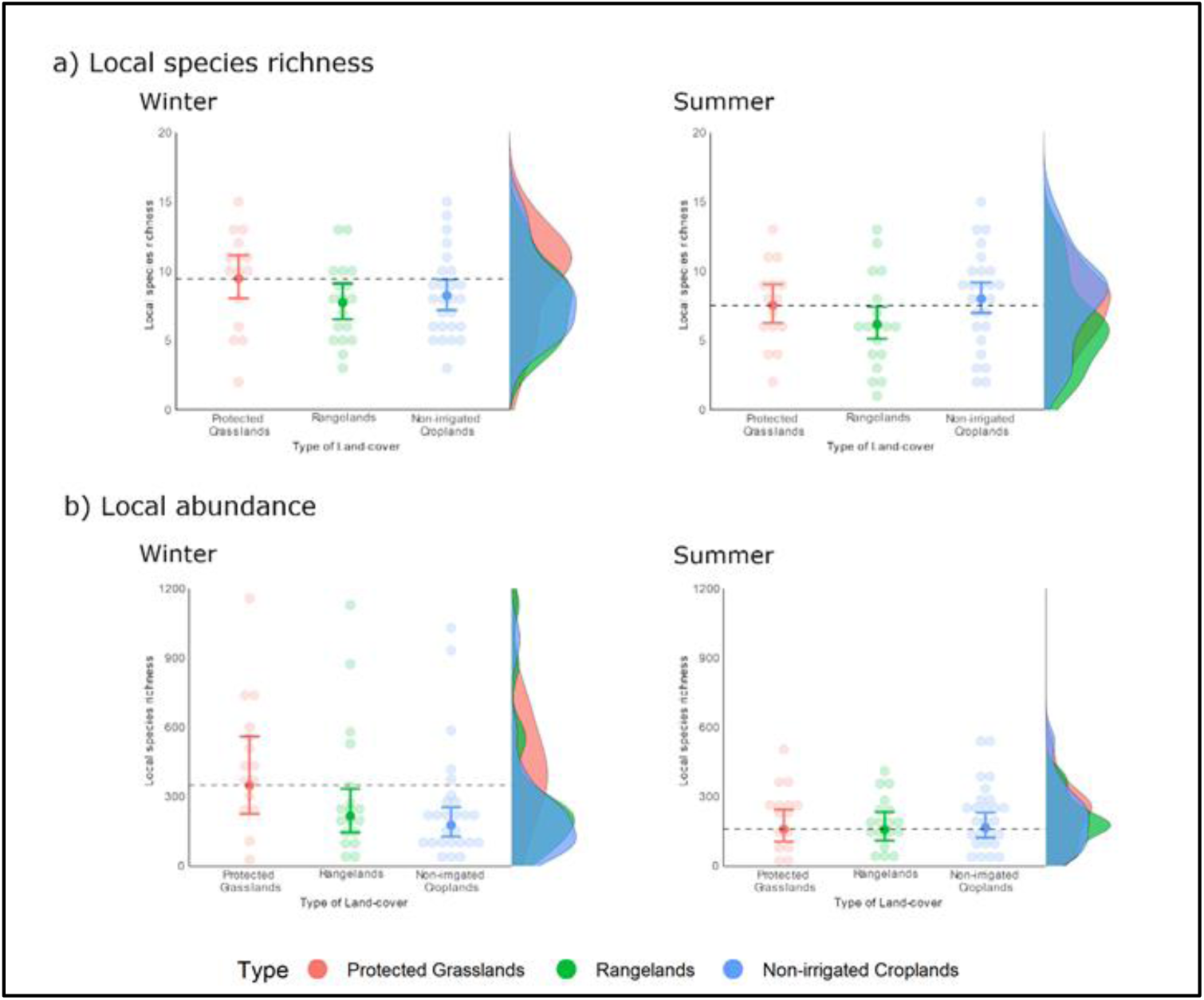
Local species richness as a function of land-cover across two seasons: a) Species richness as a function of land-cover in two seasons. b) Abundance as a function of land-cover type. In both a) and b), transparent circles indicate observed parameter value for a particular plot whereas the opaque circle represents the mean parameter value for a given land-cover type. The error bars represent 95% Poisson confidence intervals on the mean. The marginal density plot represents the distribution of parameter across sites.

#### Summer

The pattern of species richness and abundance was largely similar to winter with land-cover not explaining data any better than null models. Average species richness in protected grasslands (Mean ± SE = 7.53 ± 0.79 [6.267, 9.055]) was only marginally higher than rangelands (Mean ± SE = 6.16 ± 0.58 [5.122, 7.424]; GLM: log (Difference) = −0.2 [−0.46,0.06], p=0.13) and almost comparable to non-irrigated croplands (Mean ± SE = 8.00 ± 0.55 [6.985, 9.163]; GLM: log (Difference) = 0.06, p=0.6) (Figure 2.a.). On the other hand, average abundance in Protected Grasslands (Mean ± SE = 216 ± 33.74 [159.3, 292.8]) was not significantly different from either rangelands (Mean ± SE = 192.3 ± 30.8 [140.8, 262.7]; LM: Difference = −23.71 [−115.22, 67.79], p = 0.61) or non-irrigated croplands (Mean ± SE = 213.7 ± 25.63 [169.1, 270.1]; LM: Difference = −2.3 [−87.17, 82.56], p=0.96) (Figure 2.b.).

### 4.2 Guild structure

#### Winter

When stratified according to foraging guilds, the mean abundance of insectivores in protected grasslands was 38% higher than rangelands and 62% higher than non-irrigated croplands (Figure 3.a.). Similarly, the average abundance of granivores was 79% and 55% lower in rangelands and non-irrigated croplands, but with less precision (Figure 3.a.). The abundance of omnivores was comparable across the three land-cover types. When grouped according to habitat preference, desert and grassland specialists showed significantly lower abundance in non-irrigated croplands with a 59% decline. Mean abundance of these specialists in rangelands was 37% lower than protected grasslands, but with overlapping confidence intervals (Figure 3.b.). On the other hand, abundance of habitat generalists was highest in non-irrigated croplands followed by protected grasslands and rangelands, but with overlapping confidence intervals.

**Figure 3.**
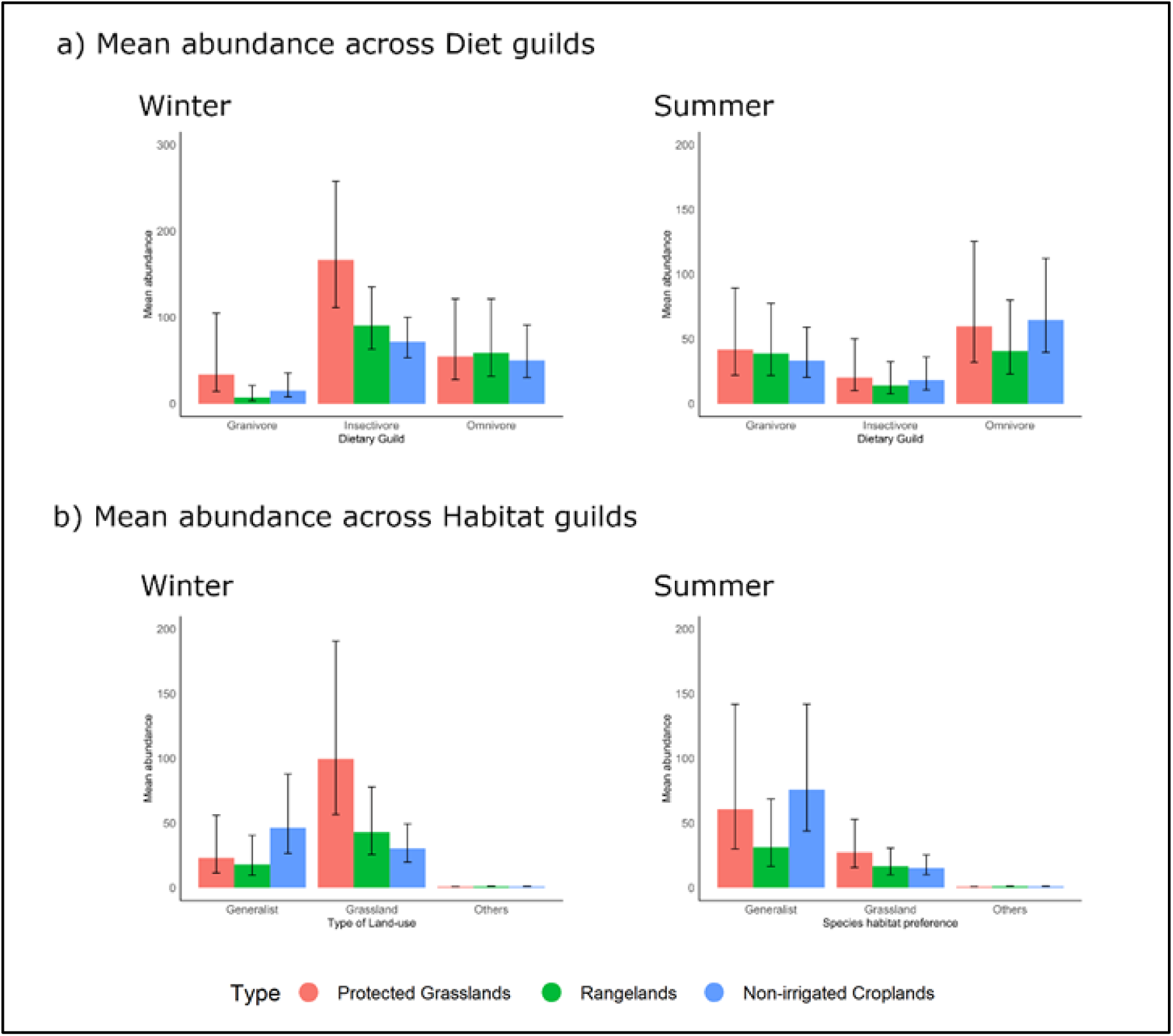
Mean abundance of birds in particular land-use (represented by the colour of the bars) stratified on the x-axis according to diet (a) and habitat (b) categories. The error bars represent 95% lognormal confidence intervals of the mean.

#### Summer

All dietary guilds and habitat guilds had comparable abundances across land-use types during the summer season (Figure 3).

### 4.3 Species pool and community composition

#### Winter

Observed and estimated species pool was highest in non-irrigated croplands and protected grasslands, followed by rangelands (Figure 4.a.). Rangelands had a significantly smaller species pool as compared to protected grasslands and non-irrigated croplands. NMDS ordination displayed weak structuring and a low compositional difference between communities of the three land-cover types (Figure 4.b.). Pairwise Multivariate Analysis of Variance test (perMANOVA) revealed that average community composition in at least one land-cover type was significantly different and land-cover type explained 6% of the total variation in community composition after controlling for the effect of location (perMANOVA: R2 = 0.06). Further inspection of results showed that during winter, community composition in rangelands and protected grasslands was only marginally different (perMANOVA: p= 0.069; ANOSIM: R=0.07). The difference between communities in protected grasslands and non-irrigated croplands (perMANOVA: p= 0.009; ANOSIM: R=0.11) was insignificant in terms of magnitude. The composition between rangelands and non-irrigated croplands was very similar (perMANOVA: p= 0.618; ANOSIM: Rr=0.02).

**Figure 4.**
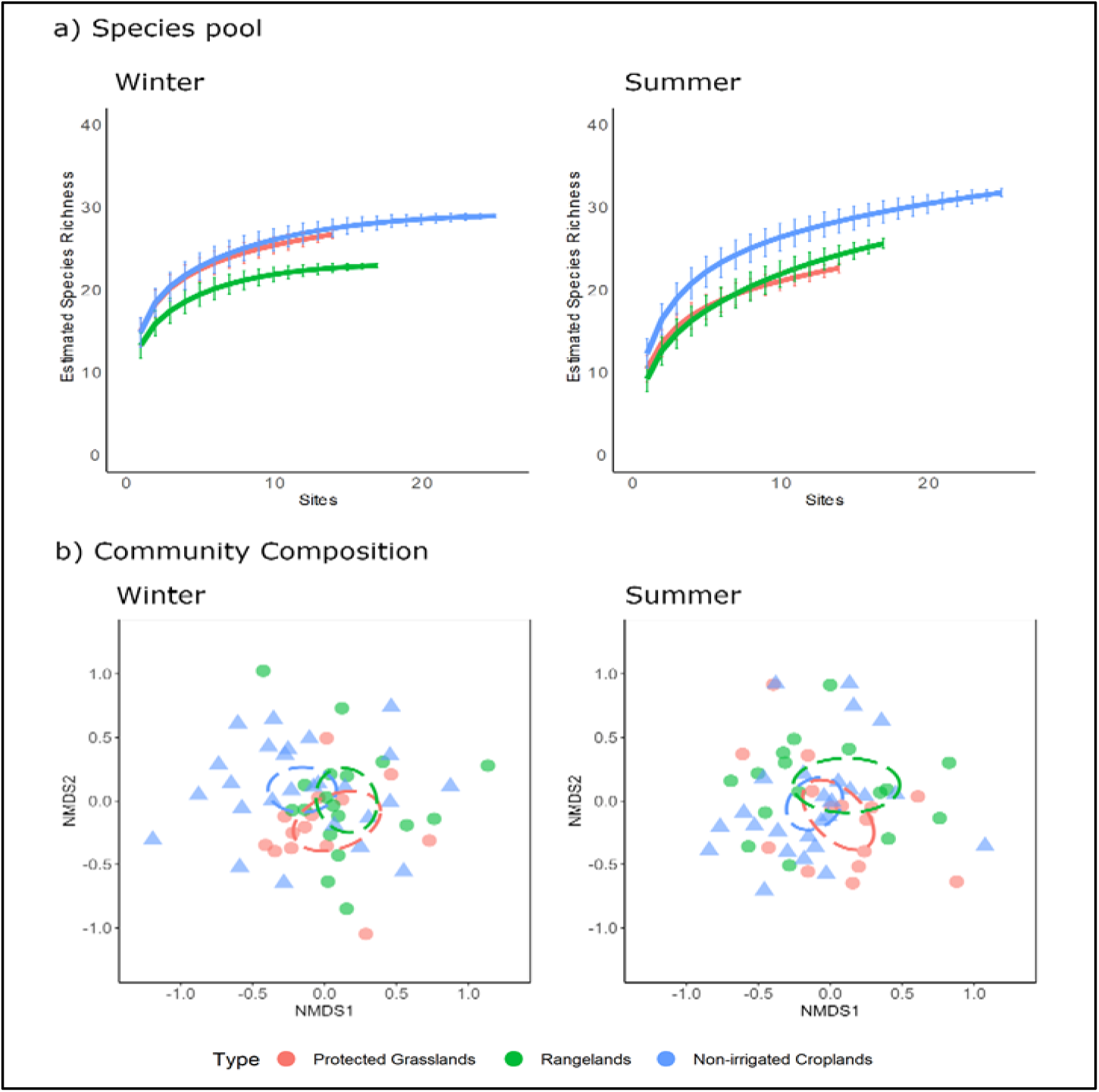
a) Sample based species-rarefaction curve for species across different land-use types, stratified according to seasons. b) NMDS ordination showing bird community composition across land-use types. Each point represents a bird community at a particular site and the colour represents the land-use type. The ellipsoids are centred at the multivariate mean of each group and the radius equals 95% confidence interval around the mean. Essentially, the ellipsoid covers the multivariate space in which the mean community composition of a particular group would lie with 95% confidence.

#### Summer

Non-irrigated croplands had a significantly larger species pool as compared to protected grasslands and rangelands (Figure 4.a.). At a local scale, land-cover could explain only 4% of the total variance in summer and no statistically significant difference was observed between the average community composition of any land-cover type (perMANOVA: R2 = 0.04). Bird communities in protected grasslands were similar to communities in rangelands (perMANOVA: p= 1.00; ANOSIM: R=0.008) and non-irrigated croplands (perMANOVA: p= 0.264; ANOSIM: R=0.03) (Figure 4.b.). Community composition in rangelands and non-irrigated croplands was also similar during both the seasons (perMANOVA: p= 1.0; ANOSIM: R=0.01).

### 4.4 Species-specific responses

#### Winter

We could derive density estimates for 20 out of the 37 species observed during the winter. When compared to protected grasslands, seven out of the twenty species (35%) had significantly lower density in non-irrigated croplands; while one species was absent and another one had significantly lower density in rangelands. Five out of seven species negatively affected by croplands and both the species negatively affected by livestock grazing were grassland specialist insectivores. On the contrary, four species in non-irrigated croplands and two species in rangelands had higher density as compared to protected grasslands. Of these, all except one species were habitat generalists and belonged to omnivorous and granivorous diet guild (Figure 5).

**Figure 5.**
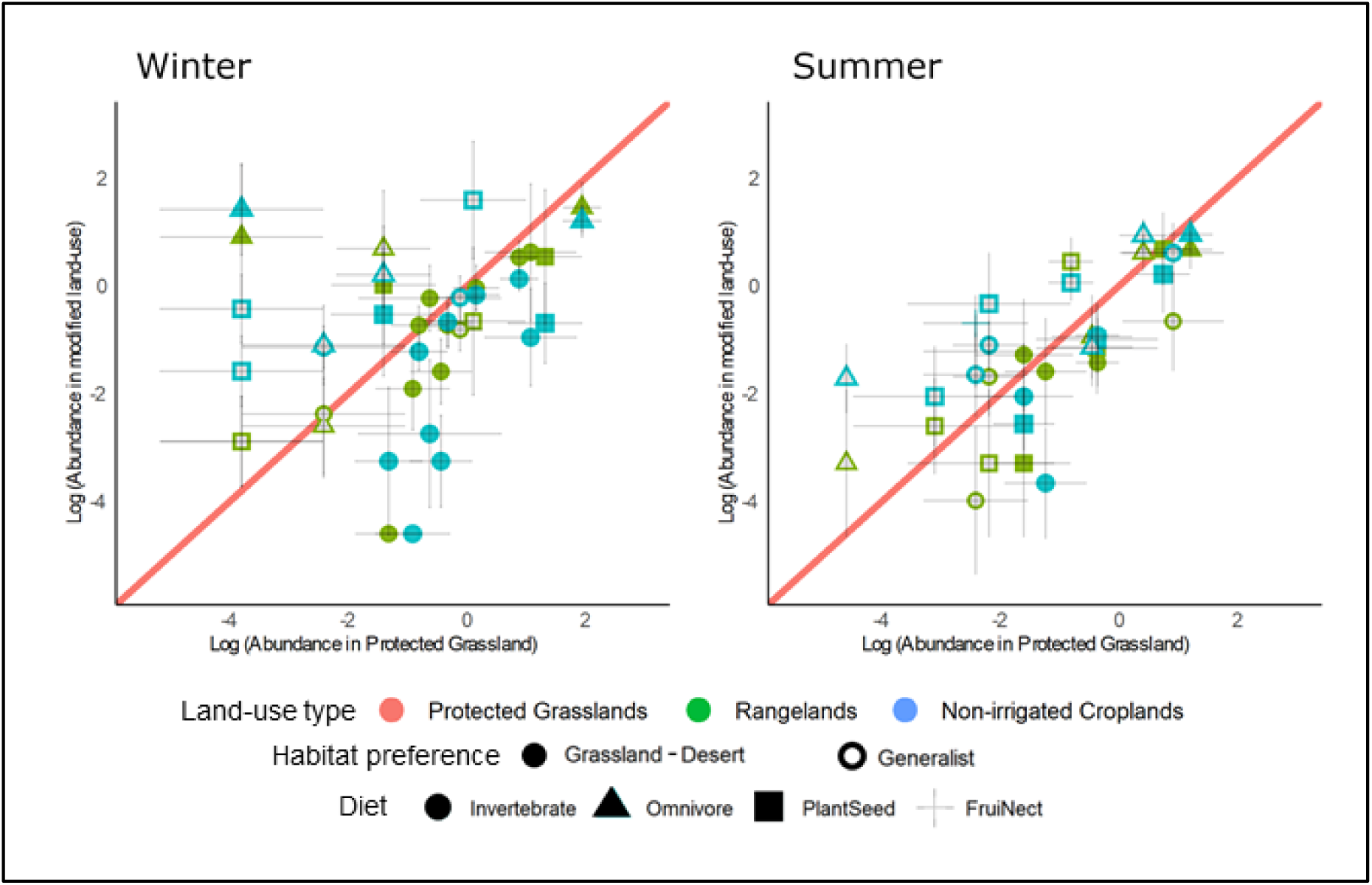
Abundance of species in a modified land-use vis-a-vis its abundance in protected grasslands (control). Points are log-transformed mean abundance of a species with y-intercept representing abundance in protected grasslands and x-intercept representing abundance in modified land-use type (represented by colour of the point). The error bars are 90% log-normal confidence intervals of the mean. The red line denotes the abundance in protected grasslands on the y-axis, such that all points above the line have higher abundance than protected grasslands (winners) and all points below have a lower abundance (losers).

#### Summer

Of the 17 species analyzed during summer, only one species (grassland specialist insectivore) had significantly lower density in rangelands and non-irrigated croplands. However, nine species in rangelands and eight species in non-irrigated croplands had insignificantly lower density as compared to protected grasslands. On the other hand, two species in rangelands and four species in non-irrigated croplands had higher density than protected grasslands. All of these winner species were habitat generalists and belonged to various habitat guilds (Figure 5).

## 5. Discussion

We examined species richness, abundance, community structure, guild structure, and individual species responses across land-use types in the Thar Desert. This is one of the first few studies to rigorously assess the impact of land-use change on arid drylands. The results are of management importance as grazing intensification, agricultural expansion, and agricultural intensification have long been identified as the main threats to the biodiversity of this landscape but their actual impacts have rarely been quantified (Singh et al., 2006). Our results indicated that near-natural protected grasslands are relatively superior in terms of sustaining avifaunal diversity and thus very important for conservation of desert avifauna, especially the specialized ecological groups within the community such as insectivores and grassland specialists. At the same time, we also found that low-intensity land-uses – rangelands and dry agriculture – harbour substantial avifaunal diversity and can complement protected areas in conserving biodiversity of this human-dominated landscape.

We found that overall local bird richness and abundance was highest in protected grasslands during both the seasons. Moreover, birds from specialized diet and habitat guilds had higher abundance in protected grasslands as compared to other land-uses. This difference in density was especially high in croplands as compared to rangelands and in winters as compared to summers. Among the species showing density declines, some were migratory Eurasian insectivores whose primary wintering ground fell in India (eg. Desert wheatear, Isabelline wheatear, Desert Warbler), while others were grassland specialist residents of India’s dryland tract (eg. Southern Grey shrike, Black-crowned sparrowlark). A common feature among all these negatively affected species was specialisation in terms of diet and/or grassland habitat preference. These specialized species are generally the ones threatened by land-use change and their conservation often revolves around protection and surgical management of habitats (Newbold et al., 2012; Sekercioglu et al., 2002). Other studies from the Thar landscape have also pointed out the importance of protected grasslands in conservation of specialised species, particularly the critically endangered Great Indian Bustard which utilizes the larger agro-pastoral landscape during the non-breeding season but is primarily dependent on good grasslands for breeding (Dutta et al., unpublished data). The needs of such specialized species and the overall higher amount of biodiversity mandates the creation of protected areas even in a relatively wildlife-friendly agro-pastoral matrix like the current one.

Contrary to popular belief and prior expectation, we found that fallow croplands and intensively grazed rangelands were also able to support a substantial amount of bird diversity in the Thar landscape. At the landscape scale, almost all species seen in protected grasslands were detected in the two modified habitat types. In fact, non-irrigated croplands had substantially larger species pool during the summer. This was due to the presence of neobiota utilising the new niches made available by agriculture and allied activities (such as the building of water tanks, sheds, etc). However, this higher species pool had little impact on the local community structure, indicating very low occupancy and abundance of the neo-colonised biota. At the local scale, bird community structure and species richness of both modified landscapes were almost identical to each other and protected grasslands. Guild and species level analyses also showed that generalist species were unaffected and in some cases, even benefited from the land-use conversion. The strikingly identical patterns during the resource-scarce summer were particularly noteworthy. These results indicate the suitability of these modified habitats for all but few species in the landscape. The major distinguishing point between the community structure of natural and modified landcover types was the abundance of diet and habitat specialist taxa, as discussed in detail above. It is worth noting, however, that specialist species (such as understory insectivores) were not completely absent in the modified landscapes but occured at significantly lower densities. These patterns are in sharp contrast to many forested systems where richness and abundance decline and community structure changes sharply in response to land-use changes (Newbold et al., 2012; Sreekar et al., 2015, but see Elsen et al., 2018; Hendershot et al., 2020;). A potential explanation for this pattern might be the very low intensity of land use, especially agriculture, in this landscape. Croplands were invariably left fallow during the dry season and thus were not very different from the other two land-uses in terms of vegetation structure. Grasslands on the other hand started losing productivity as the dry season progressed and become resource scarce by the end of winter (Dutta et al., 2017). This process was possibly accelerated in the rangelands due to simultaneous grazing pressures. All these factors result in convergence of vegetation structure of all the three land-uses by the start of summer (Kher 2019). This similarity in vegetation might be reflected in the pattern and seasonality of bird community structure across different land-covers. More research is needed to further understand the mechanisms and process driving these patterns and to find precise thresholds of land-use intensity upto which biodiversity can persist in these human modified landscapes of the Thar desert.

Taken together, these results highlight the potential utility of production land-uses (at current intensities) in supplementing protected areas for conservation of native avifaunal diversity during the resource scarce dry seasons. This is in consonance with many other studies from across the globe that have documented the suitability of low-intensity land-uses in supporting diversity of open-habitat species (Kamp et al., 2011; Wolff et al., 2001; Wright et al., 2012). Even in India, studies from other dryland systems have shown that a low-intensity agro-pastoral landscape can supplement protected areas in conservation of certain endangered bird species, especially during the dry season (Dutta, 2012; Dutta & Jhala, 2014). As these studies have focused on only one species and in semi-arid ecosystems, our multispecies study from an arid landscape increases the scope of these inferences and provides insights from an important conservation region. Finally, although we didn’t test the trade-off between biodiversity and production yields formally, our results qualitatively point towards the suitability of land-sharing approach for the management of Thar desert’s avifanual diversity. Such an approach has long been advocated for management of Indian grasslands and our results provide further evidence in its support (Anoop et al., 2017; Dutta & Jhala, 2014; Dutta, Rahmani, & Jhala, 2011; A. Rahmani, 1987; Singh et al., 2006).

Analysis like ours substitute space for time, ignoring time-lags in biological response, and thus miss out on the temporal aspects of land-use response. Similarly, these studies often ignore the role of habitat configuration in driving the observed patterns. In this study, we explicitly tested the spatial effects of change in habitat composition on bird communities but did not formally check for effects of habitat configuration on community parameters. Yet, we do not expect this to change our conclusions majorly, as most of our transects were embedded in a similar landscape matrix and the habitats were homogeneous at the scale of sampling. Another critical assumption made during such comparisons is that all human-modified areas were like present-day natural areas before the land-use intervention happened. Based on a generic literature review and discussions with senior scientists and local elders, we believe this assumption to be true in our case. A major caveat of our study is that we do not report data from the wet season (breeding season for many species). Many dryland species are known to show differential habitat preferences during the breeding and non-breeding season and data from the wet season is thus necessary before translating our conclusions to the entire year. More studies with finer scale data and longer duration are required to better understand the land-uses responses of species in the Thar desert and to effectively reconcile bird conservation with human livelihoods in this important conservation landscape.

## 6. Conclusion

Primary habitats are often considered irreplaceable for the conservation of biodiversity (Barlow et al., 2018; Lee et al., 2011) and even our results point out their relative superiority in sustaining native biodiversity. However, maintaining primary habitats is resource-intensive and in conflict with socio-economic mandates of food security, livelihood generation, and socio-economic equity (Fischer et al., 2014). This makes sustenance of biodiversity in human-modified landscapes critical from a conservation point of view. Results from this study suggest that while primary grasslands are the most suitable habitats for the birds, but livestock grazing and seasonal low-intensity farming can also support substantial amounts of avifauna during the dry season. These modified land-uses are however not conducive to all the species (eg. grassland insectivores) and during all the seasons (winter). Thus, production land-uses at current intensity seem to only supplement, but not replace, protected areas in conservation of avifauna. Evidence from this study supports the approach of conserving grasslands as large-landscapes strategically managed as agro-pastoral mosaics with small protected areas embedded within them. This strategy can potentially integrate the benefits of both land-sharing and land-sparing (Fischer et al., 2008; Kremen & Merenlender, 2018); and conserve all the regional species while not compromising on the human needs, as shown by empirical evidence gathered in this study and many others (Dutta & Jhala, 2014). However, given the single-season nature of our study, our findings are only indicative and in the future need to be augmented with multi-season species-level population data that examines how individual species’ densities depend on land-use types and habitat structure and how they relate to agricultural yields, as proposed by Green et al., 2005. Nonetheless, our results are highly relevant to the contemporary management scenario of the western Thar landscape where the highly human-dominated landscape is seeing rapid land-use change and even the sole protected area in the region is not buffered from anthropogenic change.

## Acknowledgements

VK was supported by a Master’s fellowship from the Wildlife Institute of India; and logistic support from the Bustard recovery program, Wildlife Institute of India. Fieldwork in Desert National Park WLS was permitted by the Rajasthan Forest Department under permit no. F19(13) Permission/CWLW/2017-18/1221. V.K. and S.D. would like to thank Aashi Parikh, Amedsingh, Aradeen Khan, Jogsingh, Pushkar Phansalkar, Rohit Kolhatkar and CAMPA-Bustard Recovery Program Team for assistance in fieldwork. V.K. is grateful to Navendu Page and Malvika Onial for supervising parts of the study; and YV Jhala, Q Qureshi, Gopi GV and GS Rawat for guidance and support. V.K. is also thankful to his thesis examination committee, especially Suhel Qader, for critical comments and suggestions.

## Notes

### Competing Interest Statement

The authors have declared no competing interest.

